# Predicting Immunogenic CD4+ T Cell Epitopes in Bacteria Using Antigen and Peptide Features

**DOI:** 10.1101/2024.10.25.620357

**Authors:** Daniel Marrama, Hannah Battey, Ehdieh Khaledian, Miriam Müller, Sudhasini Panda, Ricardo da Silva Antunes, Alessandro Sette, Cecilia S. Lindestam Arlehamn, Bjoern Peters

## Abstract

**Background:** T cell epitope prediction methods have been broadly utilized to facilitate epitope discovery in infectious agents and help design reagents, diagnostics, and vaccines. Current prediction methods are mainly focused on peptide presentation by MHC molecules, which is a necessary but not sufficient requirement for an epitope. For complex pathogens such as bacteria, it would be desirable to make such predictions more specific to limit the number of candidates that have to be experimentally tested.

**Objective:** To develop a machine learning-based prediction model that integrates both peptide-level and antigen-level features to improve the specificity of CD4+ T cell epitope predictions for bacteria.

**Methods:** We used a dataset of 20,216 peptides from *Mycobacterium tuberculosis* (Mtb), tested for T cell recognition in Mtb-infected participants, that led to the discovery of n = 144 peptide epitopes. For each peptide, we calculated six peptide-level features (e.g. MHC class II binding predictions and conservation scores) and six antigen-level features (e.g. including RNA expression levels and subcellular localization scores). Three machine learning algorithms—Random Forest, Gradient Boosting, and XGBoost—were trained using stratified, 5-fold cross-validation and combined into an ensemble model. Experimental validation was performed on *Streptococcus pneumoniae* peptides, using ex vivo IFNγ assays to confirm the predictive performance.

**Results:** The ensemble model achieved an ROC-AUC of 0.91 in predicting immunogenic peptides in the *Mycobacterium tuberculosis* (Mtb) dataset. Gene expression and conservation were identified as the most impactful features, followed by MHC class II binding predictions. When validated on an independent *Bordetella pertussis* dataset, the model demonstrated accurate predictive capability, especially for peptides with broad recognition in the participant cohort (ROC-AUC up to 0.82). Prospectively applying the model to *Streptococcus pneumoniae*, we synthesized peptides predicted by our ensemble model to be immunogenic or non-immunogenic. Ex vivo testing with PBMCs from healthy participants showed that peptides predicted to be immunogenic elicited significantly higher IFNγ responses than non-immunogenic peptides, validating the model.

**Conclusions:** Our machine learning approach, integrating both peptide and antigen features, effectively predicts immunogenic CD4+ T cell epitopes across different bacterial pathogens. This method enhances epitope selection efficiency, aiding vaccine development and immunological research by reducing the need for extensive experimental screening.

## 1. Introduction

T cell lymphocytes recognize linear peptides, or epitopes, presented to them via major histocompatibility complex (MHC) molecules on host cells. CD4+ T cells recognize epitopes presented by MHC class II molecules, which are predominantly expressed on professional antigen-presenting cells, such as macrophages. These cells are the primary source of T cell epitopes derived from bacteria (1).

T cell epitope predictions have been successfully used to identify candidate peptides from infectious agents, allergens, and cancer cells (2). These predictions have predominantly relied on the features of the peptides themselves, especially their predicted MHC binding affinity. When the goal is to identify peptides likely to be immunogenic in the general population, predictions of MHC class II binding can narrow down the number of peptide candidates to the top ∼20% (3). However, in the case of complex bacterial pathogens expressing thousands of antigens, this will still leave tens-to hundreds of thousands of possible candidate peptides.

To address this problem, we investigated the development of prediction methods that integrate the properties of specific peptides, such as their binding affinity, with the properties of the antigens that they are derived from. For example, we and others have shown that the expression level of an antigen impacts the likelihood that peptides contained in it will be recognized by T cells (4–6). Additionally, we found that certain antigen properties can impact their immunogenicity, such as proteins secreted by or contained within type 7 secretion systems (T7SS) being the main target of immune responses in individuals who are IGRA+ (7). At the peptide level, we observed that beyond MHC binding, the degree of conservation of a peptide in related antigens can increase the likelihood of T cell recognition while conservation in the host can reduce it (8,9). We set out to combine these findings into a unified prediction method for bacterial pathogens using machine learning approaches.

In this study, we describe the peptide prediction model features and report the contribution of each feature using an existing Mtb dataset for training (7). Next, we assessed the versatility of the prediction model by applying it to an existing *Bordetella pertussis* dataset. To further demonstrate the model’s ability to successfully discriminate between immunogenic and non-immunogenic peptides, we synthesized peptides with the highest and lowest predicted values and tested for immunogenicity *ex vivo* in PBMCs from healthy individuals. We validated our model experimentally using *Streptococcus pneumoniae*, the most common cause of bacterial pneumonia (10), and PBMCs from healthy individuals. Considering that most of the general population has been exposed to *S. pneumoniae* by adulthood (10), we used PBMCs from healthy controls to screen for immune responses against synthesized peptides which were predicted either immunogenic or non-immunogenic for comparison. Finally, we combined the Mtb and *Bordetella pertussis* datasets to train a combined model to be used for broader CD4+ T cell epitope prediction against bacterial pathogens.

## 2. Results

### 2.1 Model Development and Cross-Validation on a Mtb Dataset

We utilized a dataset of 20,216 peptides derived from *Mtb* that was systematically tested for T cell recognition in Mtb-infected participants (IGRA+; see Methods and (7)). That peptide set was created to represent each Mtb protein with 2-10 peptides and prioritized peptides in each protein that had highly predicted promiscuous MHC class II binding based on the methods available. A total of 144 peptides were recognized by at least two participants, and these were considered positive labels in training our machine learning models, while the remainder of the peptides were considered negatives.

For each peptide in the set, we calculated six features on the peptide level, namely MHC II binding predictions (7-allele score (3)), Mtb strain conservation scores (allowing 0-3 amino acid substitutions with one score for each threshold), and the best peptide match to the human proteome. We also calculated six features on the protein level, namely RNA expression as TPM values, subcellular localization prediction scores (a score for each component: cytoplasm, cytoplasmic membrane, outer membrane, and extracellular) using PSORTb (11), and protein existence levels as curated by UniProt (**Table 1**). These features were selected based on prior publications showing their importance in immune recognition (4–9). Additionally, we added a random value between 0 and 1 to each peptide to determine its ranking in feature importance.

**Table 1.** Features collected for Mtb peptides at the peptide level and antigen level.

| Level | Feature | Method/Source |
| --- | --- | --- |
| Peptide | MHC-Binding 7-allele score | NetMHCIIpan |
| Peptide | Mtb strain conservation (exact match) | PEPMatch |
| Peptide | Mtb strain conservation (1 substitution) | PEPMatch |
| Peptide | Mtb strain conservation (2 substitutions) | PEPMatch |
| Peptide | Mtb strain conservation (3 substitutions) | PEPMatch |
| Peptide | Best human homology percentage | PEPMatch |
| Peptide | Random Control Value (for feature importance) | Python `random` library |
| Protein | RNA Expression (TPM) | GEO Dataset |
| Protein | Cytoplasm localization score | PSORTb |
| Protein | Cytoplasmic membrane localization score | PSORTb |
| Protein | Outer membrane localization score | PSORTb |
| Protein | Extracellular localization score | PSORTb |
| Protein | Protein Existence Level | UniProt |

We applied three different machine learning algorithms: XGBoost, Gradient Boosting, and Random Forest. Each algorithm was used to train a model using grid search hyperparameter tuning with stratified 5-fold cross-validation due to class imbalance. Model performance was evaluated using the area under the curve (AUC) for the Receiver Operating Characteristic (ROC) and AUC0.1, the AUC between 0 and 0.1 false positive rate. When evaluating performance on the test data, the Random Forest model achieved an AUC of 0.88 and an AUC0.1 of 0.81. The Gradient Boosting model yielded an AUC of 0.85 and an AUC0.1 of 0.79, while the XGBoost model demonstrated an AUC of 0.90 and an AUC0.1 of 0.78. The ensemble model, which combined predictions from all three methods by averaging probabilities, further improved predictive performance, achieving an AUC of 0.91 and an AUC0.1 of 0.82. The ROC curves with AUC values of each model on the test data are shown in **Figure 1**.

**Figure 1.**
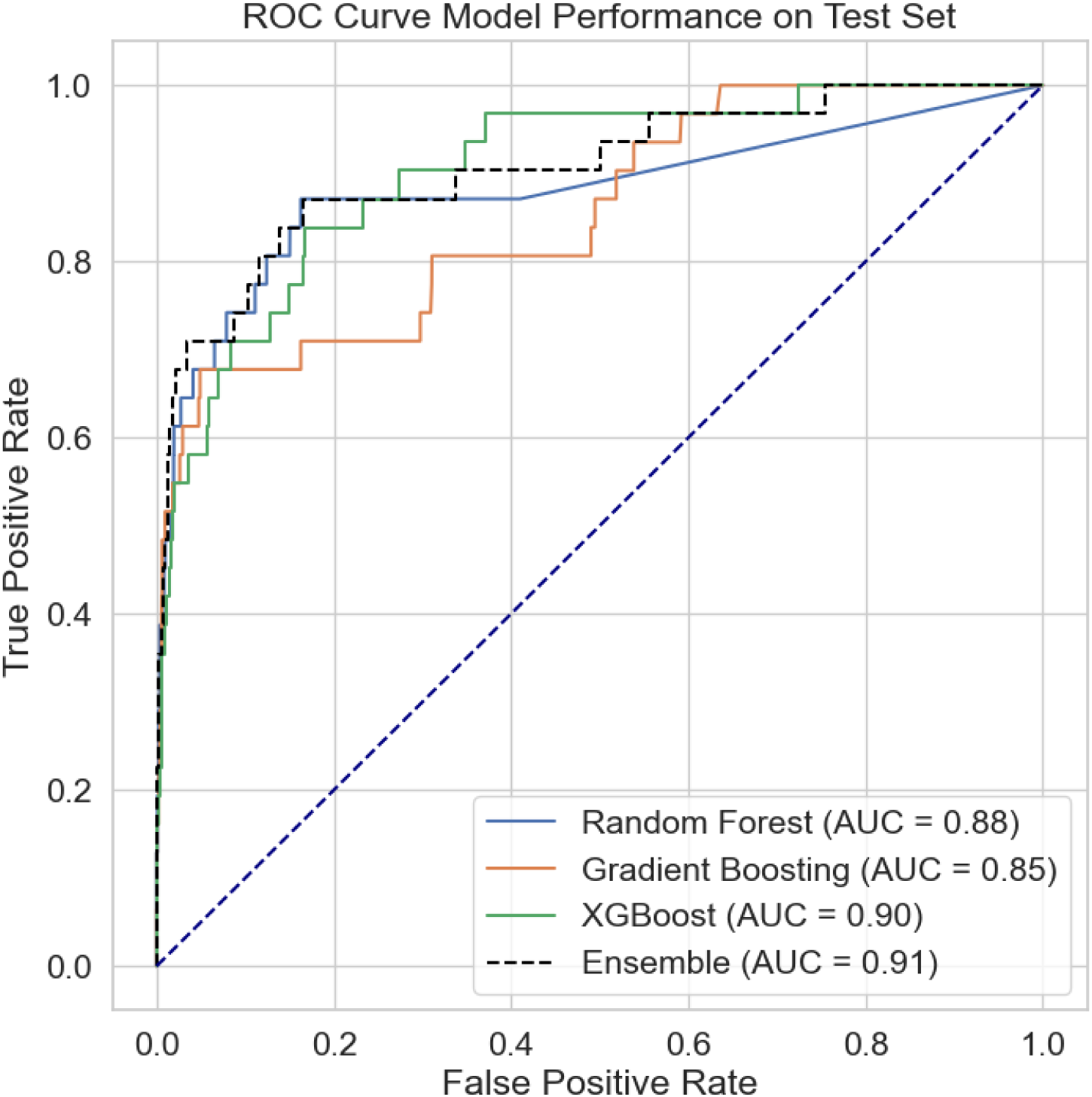
Performance of immunogenicity prediction by the trained models on the withheld testing data using ROC curves. Performance was evaluated as ROC-AUC for each model trained (Random Forest, Gradient Boosting, and XGBoost) and the ensemble of the three models. The ensemble model achieved the highest AUC of 0.91, outperforming each individual model.

### 2.2 Feature Importance Analysis

To determine how different features contribute to the predictive performance of the model, we implemented a permutation approach where each feature has its values randomly shuffled one at a time, and the decrease in model performance as measured by ROC-AUC was assessed. This method was applied individually to each of the three algorithms: Random Forest, Gradient Boosting, and XGBoost, and the results for each feature were aggregated (**Figure 2A**).

**Figure 2.**
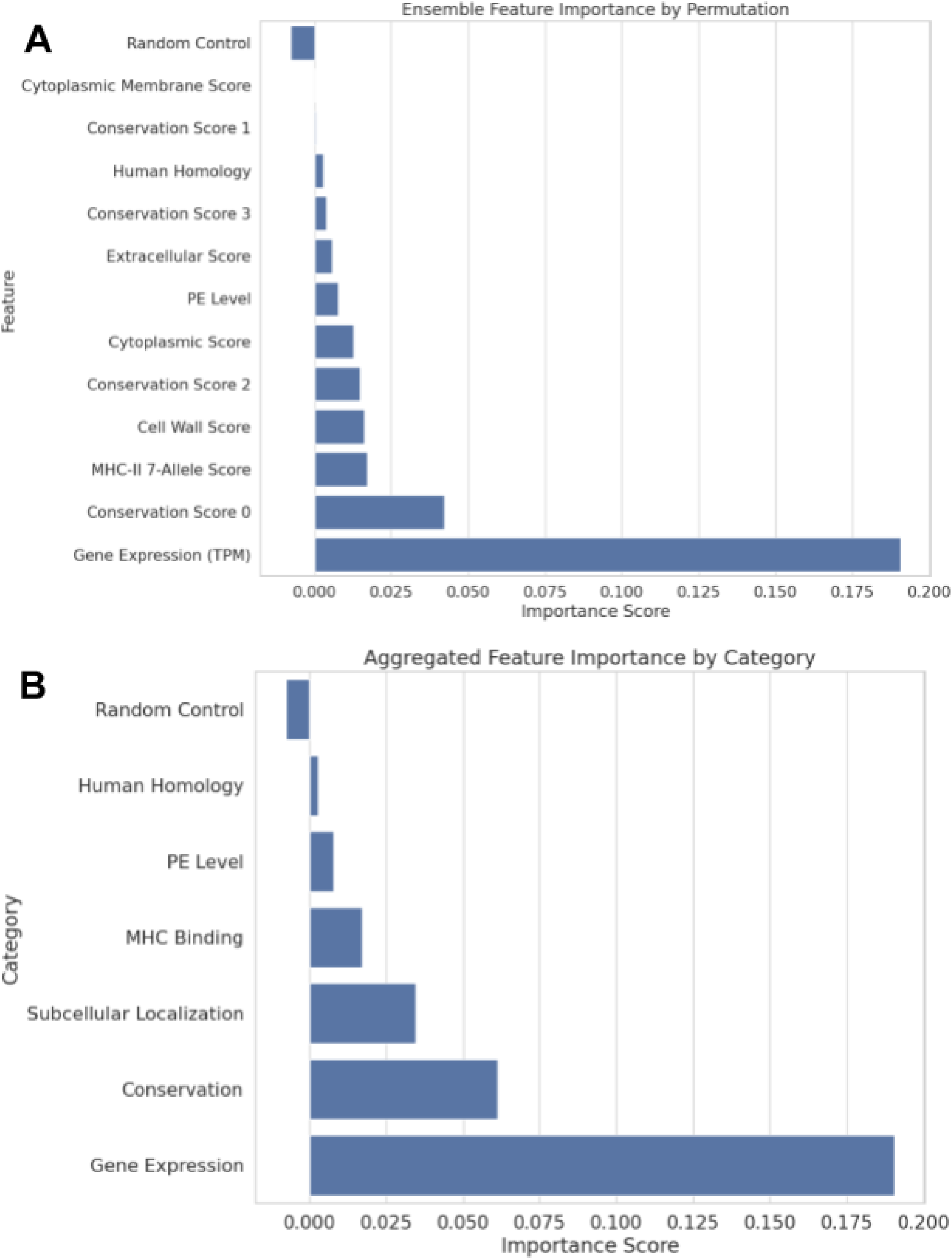
Feature importance scores from the ensemble model as assessed by permutation after training. A) Each feature used for the training of each machine learning model is evaluated by the loss of ROC-AUC performance when the feature’s values were permuted. Higher bars indicate greater importance to the model’s predictive ability. B) Feature categories were combined to show the overall contribution of broader groups, highlighting which biological factors had the most influence on model performance, with gene expression having the greatest impact, followed by conservation and subcellular localization.

By far, the most impactful feature was the gene expression associated with each peptide antigen. Other features, such as conservation (exact match across the 15-mer peptide and allowing 2 amino acid substitutions), MHC-II binding prediction, and the cell wall score from PSORTB, also demonstrated an effect on performance. Conversely, certain features like cytoplasmic membrane score and best peptide match to the human proteome had minimal impact. The random control feature scored the lowest, indicating that all the rest of the collected features had some positive effect on the model performance, even if the effect was minimal.

A disadvantage of the permutation approach for the individual feature importance evaluation is that correlated features such as the conservation scores 0, 1, 2, and 3 and the different subcellular localization scores cannot be properly evaluated, as leaving one such feature out will allow the model to compensate for it with the information in others. We thus also combined all features from the same category and left them out in a group for the permutation to assess their overall contribution to model performance (**Figure 2B**). This showed that gene expression level was the leading feature, followed by the combined conservation features and the combined subcellular localization features. Given that the peptides tested in the Mtb dataset were all pre-selected for their high MHC binding capabilities, the ability of such scores to discriminate epitopes from non-epitopes cannot be fully assessed.

### 2.3 Model Performance on B. pertussis Data

We set out to test if the model trained on the Mtb peptide dataset would have predictive power for a completely independent pathogen: *Bordetella pertussis,* the causative agent of whooping cough. We utilized a dataset of 24,294 peptides derived from *Bordetella pertussis,* which had been screened in 20 participants (as described in Methods and (12)). Given the variance in performance of the model based on the positivity threshold we described above, we assessed model predictions using ROC-AUC using variable thresholds ranging from one to four participants.

At the threshold of recognition by at least one participant, the ensemble model was tested against 2,939 positive peptides, yielding an AUC of 0.50. As the participant recognition threshold increased to two, the model was evaluated with 412 positive peptides and achieved an AUC of 0.58. With peptides recognized by at least three participants, the model’s performance was tested against 79 peptides, resulting in an AUC of 0.78. This pattern of assessment continued as the threshold increased, with the highest AUC achieved of 0.87 for the 32 peptides recognized by at least four participants. The trend observed in the model’s performance on the *Bordetella pertussis* dataset shows the model is more adept at identifying peptides with broad recognition among the participant population as opposed to peptides only recognized at the individual level. The ROC curves and AUC values of the ensemble model prediction at each participant threshold are shown in **Figure 4**.

**Figure 4.**
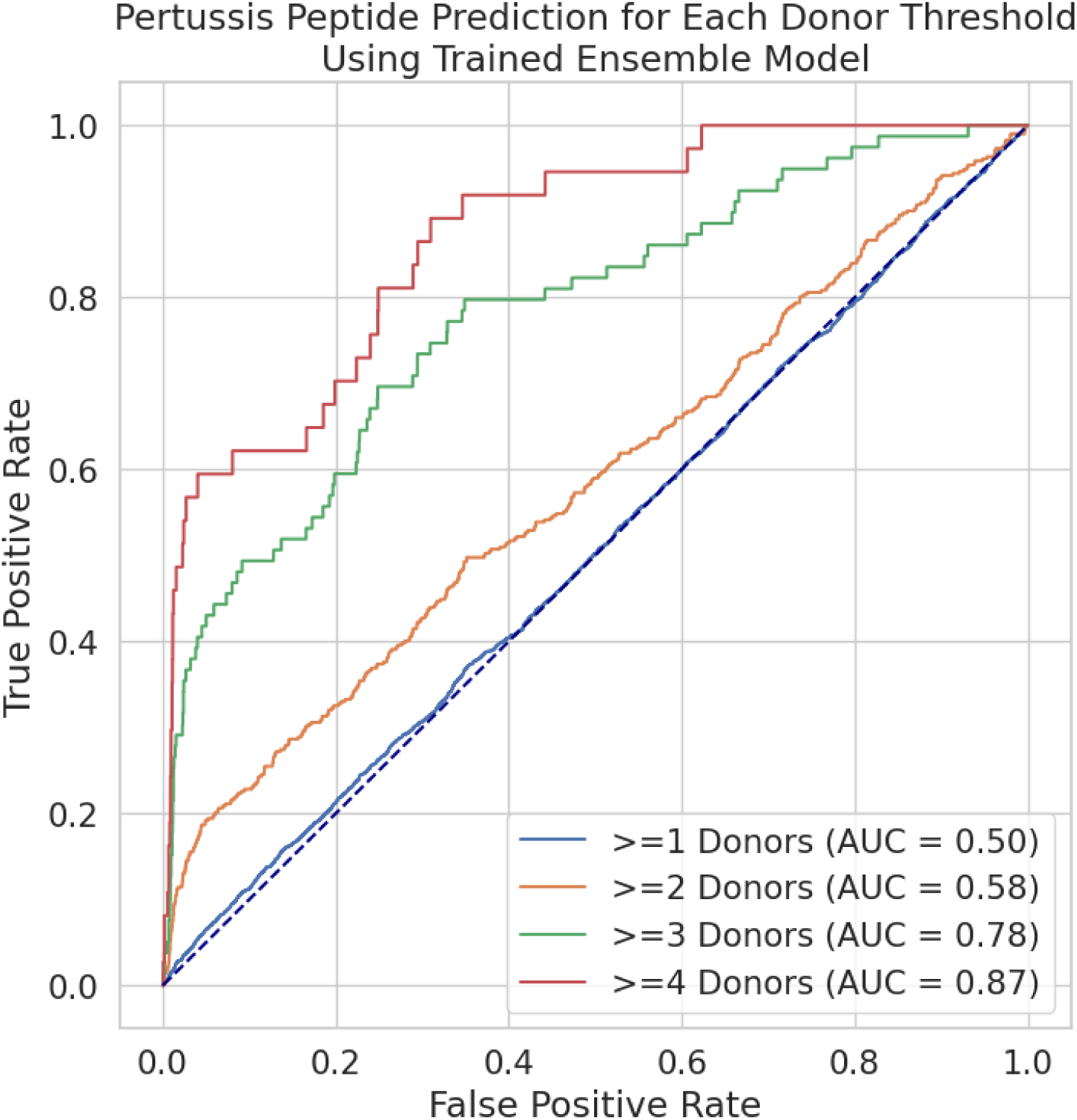
Performance of ensemble model predicting *Bordetella pertussis* epitopes at various donor thresholds. The model progressively improves performance as the participant recognition threshold increases, with the highest AUC of 0.87 achieved at the four-participant threshold. This suggests the model is better at predicting peptides with broader recognition across multiple participants.

### 2.4 Model Performance to Identify Immunogenic S. pneumoniae Peptides in a Prospective Fashion

Next, we wanted to test, in a prospective fashion, if the methods we have developed can identify immunogenic peptides from an independent pathogen when starting from the entire proteome better than when using MHC presentation predictions alone. We chose *S. pneumoniae* as a target since it is the most common cause of community-acquired pneumonia (CAP), and most adults have been exposed to the pathogen through infection or colonization during their lifetime (10). We wanted to specifically test the ability of our model to select peptides that are immunogenic from *S. pneumoniae* over what would be selected based on MHC binding predictions alone. The peptides with the highest and lowest ensemble probability scores were classified into one of two groups for peptide synthesis: predicted to be immunogenic (n=130) and predicted to be non-immunogenic (n=130). As expected, the predicted immunogenic peptides had lower (=better) protein existence scores and higher conservation, gene expression, and MHC-II binding scores than those with low predicted immunogenicity (**Figure 5**). Despite taking MHC-II binding into account prior to prediction to maximize the impact of the other features for peptide selection, peptides with higher ensemble probability scores achieved higher MHC-II scores.

**Figure 5:**
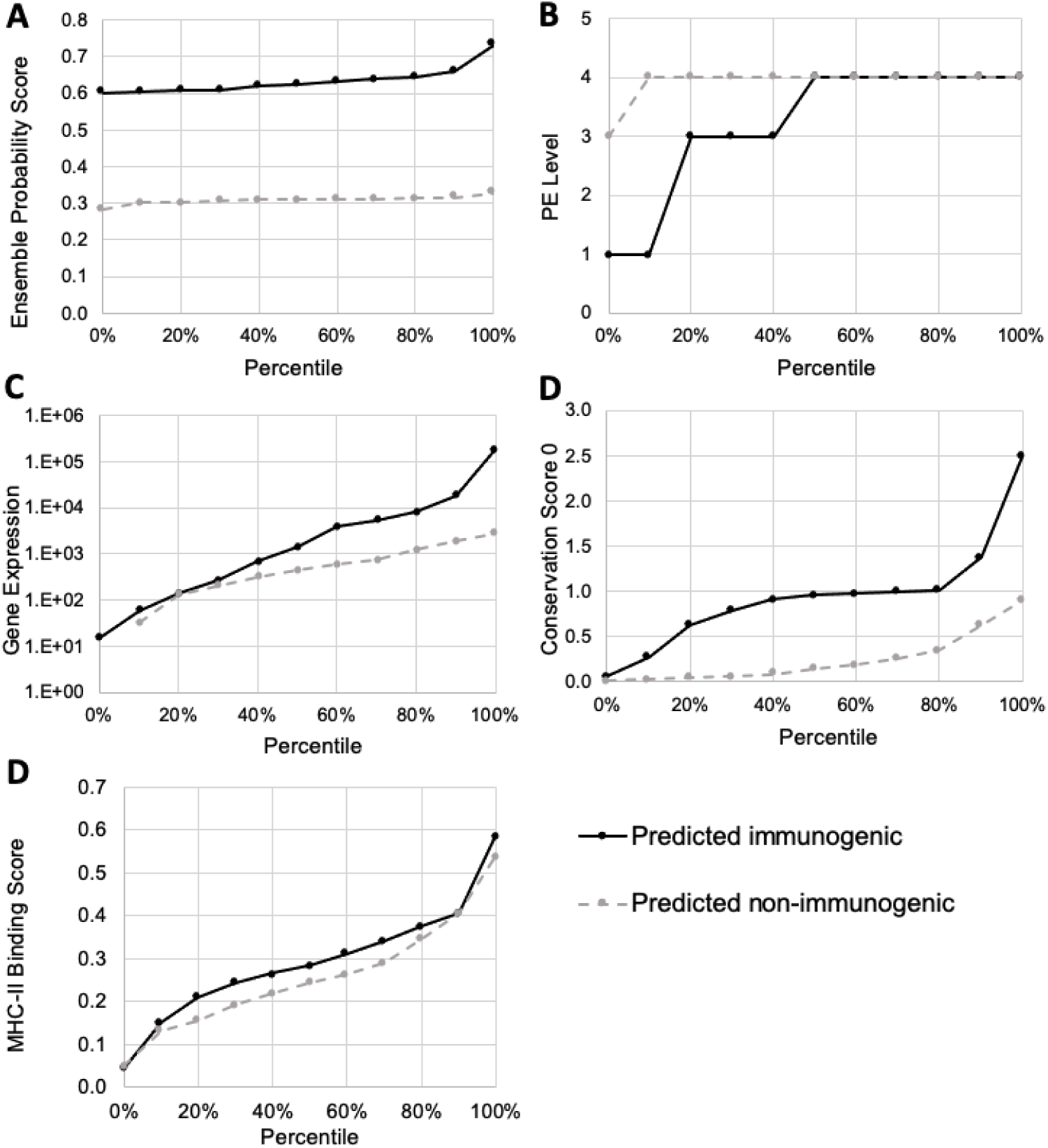
Overview of *Streptococcus pneumoniae* feature characteristics among peptides predicted to be immunogenic and non-immunogenic using the ensemble model. A) The highest (n=130) and lowest (n=130) *S. pneumoniae* peptide probability scores were used to select ‘predicted immunogenic’ and ‘predicted non-immunogenic peptides’. As expected, peptides predicted to be immunogenic had B) lower protein existence levels (which means better evidence for existence), C) higher gene expression, D) higher conservation across strains (no amino acid substitutions), E) and higher predicted MHC-II binding compared to peptides with a low ensemble probability score compared to peptides predicted to be non-immunogenic based on the ensemble model.

The 130 highest and 130 lowest probability-scored peptides were synthesized for screening in PBMCs from healthy participants (n=20). Peptides were combined into 26 pools of 10 peptides (P1-P26) each to minimize the number of cells required for initial screening. IFNγ, one of the most prevalent pro-inflammatory cytokines, was measured using Fluorospot assay following stimulation with peptide pools, and all responses were recorded. The peptide pools were tested alongside a megapool (MP) containing all 260 peptides, a negative control, and a positive control (**Figure 6A**). Predicted immunogenic pools (P1-P13) had a higher magnitude of IFNγ responses compared to predicted non-immunogenic pools (P14-P26) (p=0.003). The megapool and a single 10-peptide pool (P7) were capable of eliciting a strong IFNγ response.

**Figure 6:**
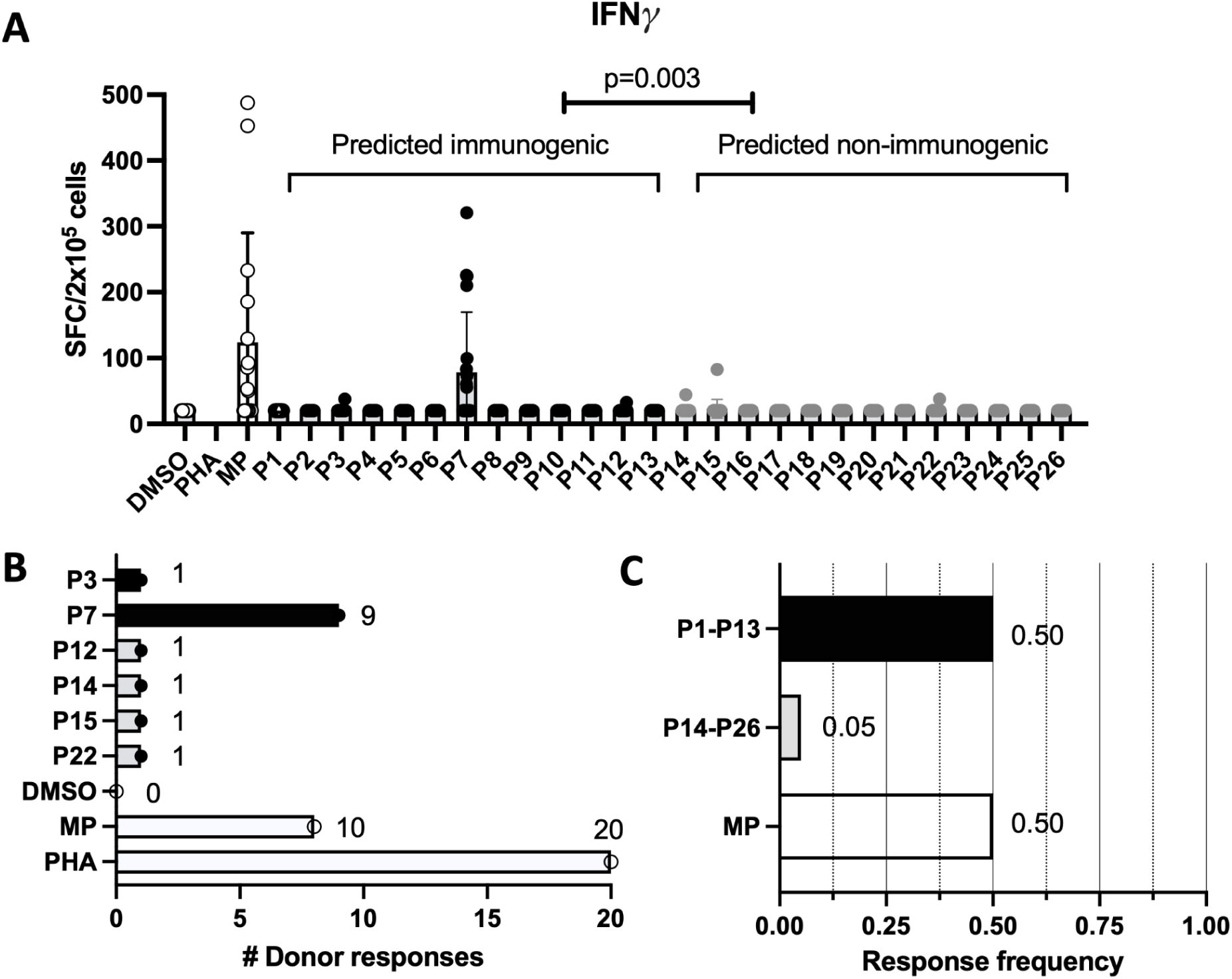
Peptides with high predicted immunogenic potential elicit a higher magnitude IFNγ response across a higher frequency of participants. A) Cytokine production was measured by IFNγ Fluorospot following stimulation with DMSO (negative control), PHA (positive control), megapool containing all 260 peptides (MP) and 10-peptide pools (P1-P26). Each data point represents the number of SFC per well (2×10^5^ cells) for each participant (n=20), with bars indicating means with standard deviations. The magnitude of IFNγ SFC per well for immunogenic versus non-immunogenic peptides was compared with an independent one-tailed t-test. Participant responses to stimuli were considered positive if the following 3 criteria were met: 1) the number of SFC above the background was >20, 2) a greater than 2-fold increase above the background, 3) p < 0.05 by independent one-tailed t-test and Poisson distribution test when comparing stimuli to negative control. A maximum cutoff of 500 SFC/2×10^5^ was utilized. Any responses that did not meet the 3 criteria for positivity were considered IFNγ negative and plotted as 0 SFC/2×10^5^ cells. B) The number of participants with a positive IFNγ response following stimulation. Each dot represents the average SFC across triplicates. C) The response frequency was calculated by dividing the number of participants with a response by the total number of participants screened. The peptide pools were tested as individual 10-peptide pools and aggregated based on any response to P1-P13 or P14-P26. The megapool was tested as a single pool of 260 peptides.

Out of the 20 participants, P7 elicited a positive IFNγ signal in 9, compared to single-participant responses in P3, P12, P14, P15, and P22 (**Figure 6B**). The megapool was recognized by 10/20 (50%) participants, with the majority coming from the predicted immunogenic peptides, suggesting that this comparably small number of 260 peptides already gives a good response compared to e.g. the >20,000 tested for Mtb.

To further evaluate the ability of the prediction model to discriminate between immunogenic and non-immunogenic peptides, we can compare the overall response frequency of P1-P13 and P14-P26 (**Figure 6C**). Half of all participants responded to at least one of the predicted immunogenic pools (P1-P13), compared to a 5% response frequency (single responder) among the predicted non-immunogenic pools. Furthermore, we can accomplish the same response frequency (50%) with half the amount of peptides (n=130) by restricting to those with the highest ensemble probability scores. The ensemble prediction model successfully identified immunogenic peptides, showing high response frequencies among the top-scoring peptides, demonstrating its potential for efficient peptide selection.

Given the dominant response to the P7 pool, we wanted to determine which specific peptides are targeted by measuring the response against each of the 10 contained peptides via IFNγ Fluorospot (**Figure 7A**). Three individual peptides were recognized among participants that responded to P7, labeled as (peptide) 1, 5, and 6. Peptide 1 was recognized in 8/9 participants, and peptides 5 and 6 were recognized in 1/9 and 6/9 participants, respectively. The start and stop indexes of peptide 1 (185/200) and 6 (184/199) are nearly identical and exhibit high sequence homology (73%, 11/15). Notably, peptides 1 and 6 share identical ensemble probability scores, protein existence levels, and MHC-II expression scores, while peptide 6 had higher conservation (exact match) and gene expression values, respectively (**Figure 7C**).

**Figure 7:**
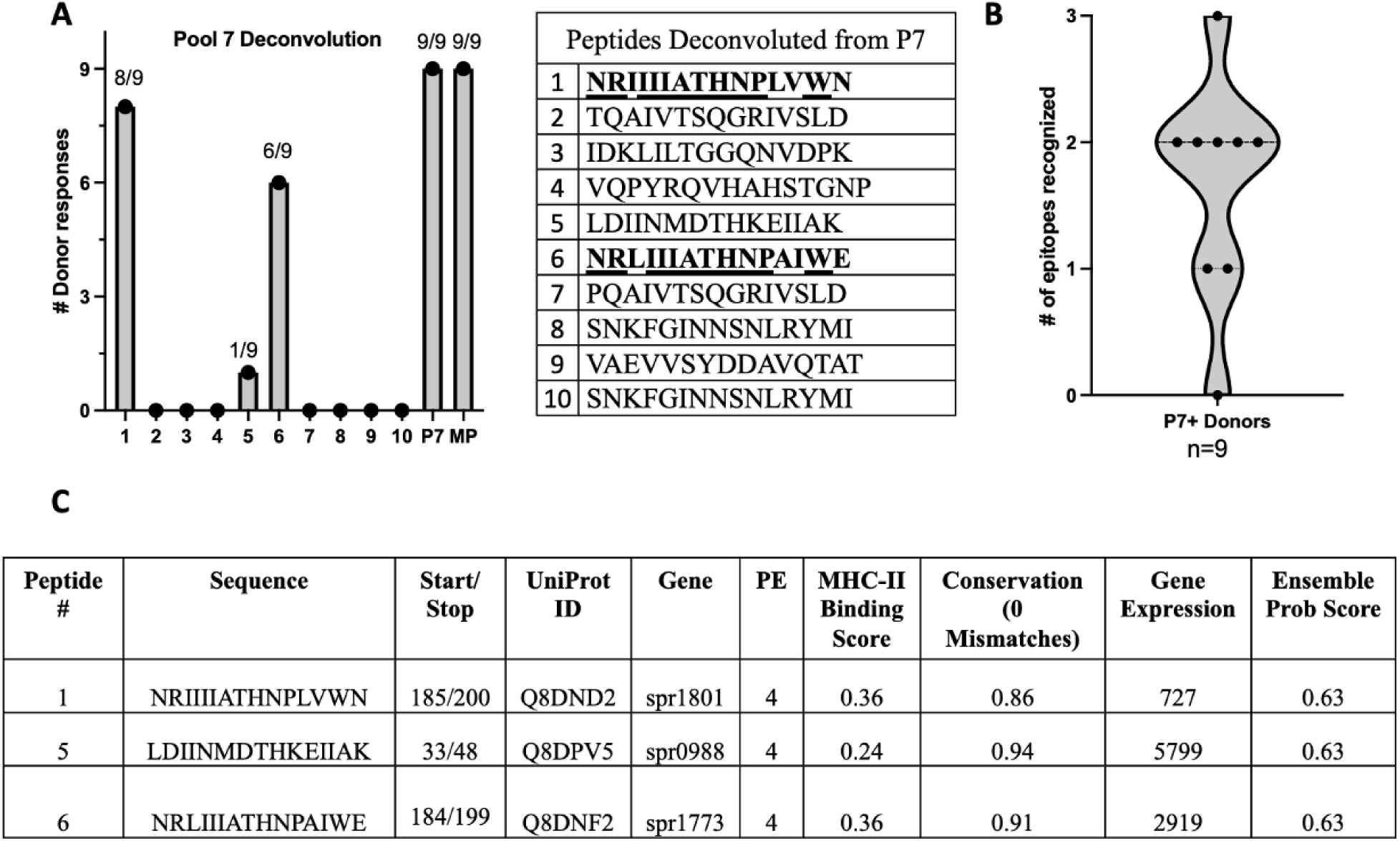
Deconvolution of P7 into its 10 peptide components reveals two T cell epitopes with similar features and sequences. A) The number of participants with a positive IFNγ response following stimulation with their respective peptides (1–10), P7, and the megapool (MP). The peptide sequences corresponding to peptides 1-10 are provided, with peptides 1 and 6 **bolded** to highlight their high performance (6/9 participant responses) and <u>underlined</u> to highlight high sequence similarity (73%). B) The number of epitopes recognized by the P7-positive participants (n=9) are visualized with a violin plot. C) Several ensemble model prediction features (MHC-II binding, conservation, and gene expression) are compared for peptides 1, 5, and 6, along with their probability scores and UniProt specifications (Start/Stop, UniprotID, Gene).

Although encoded in different genes, the proteins they are derived from are both “ABC transporter domain-containing” proteins. Furthermore, these proteins (UniProt IDs: Q8DND2 and Q8DNF2) share the same domain ID from ProSite (13) (PS50893) for the domain where the peptides are found. This deconvolution methodology illustrates that our model can be applied translationally for bacterial epitope discovery.

### 2.5 Combined Mtb and B. pertussis Model

Since we have demonstrated the predictive capabilities of our machine learning models on individual bacterial datasets, we next trained a unified model on the combined data from Mtb and *B. pertussis*. This is aimed at providing a generalized prediction tool capable of identifying immunogenic CD4+ T cell epitopes across different bacterial species, thus enhancing the utility of our method for broad-spectrum vaccine development.

The datasets from both Mtb and *Bordetella pertussis* combined to comprise 44,510 peptides—20,216 from Mtb and 24,294 from *B. pertussis*. Each peptide was annotated with the same features as in the individual models, ensuring consistency across the combined dataset.

Positive peptides were defined as those recognized by at least two participants in their respective cohorts, yielding a total of 556 positive peptides. The Random Forest, Gradient Boosting, and XGBoost algorithms were retrained on this combined dataset using the same hyperparameter tuning and cross-validation procedures outlined previously.

The ensemble model, created by averaging the probabilities from the three algorithms, achieved an ROC-AUC of 0.83 and AUC0.1 of 0.72 on the test data (**Figure 8A**). Feature importance analysis revealed that gene expression still had the greatest impact on the model prediction, but conservation and subcellular localization had more impact than the original Mtb model did (**Figure 8B**).

**Figure 8.**
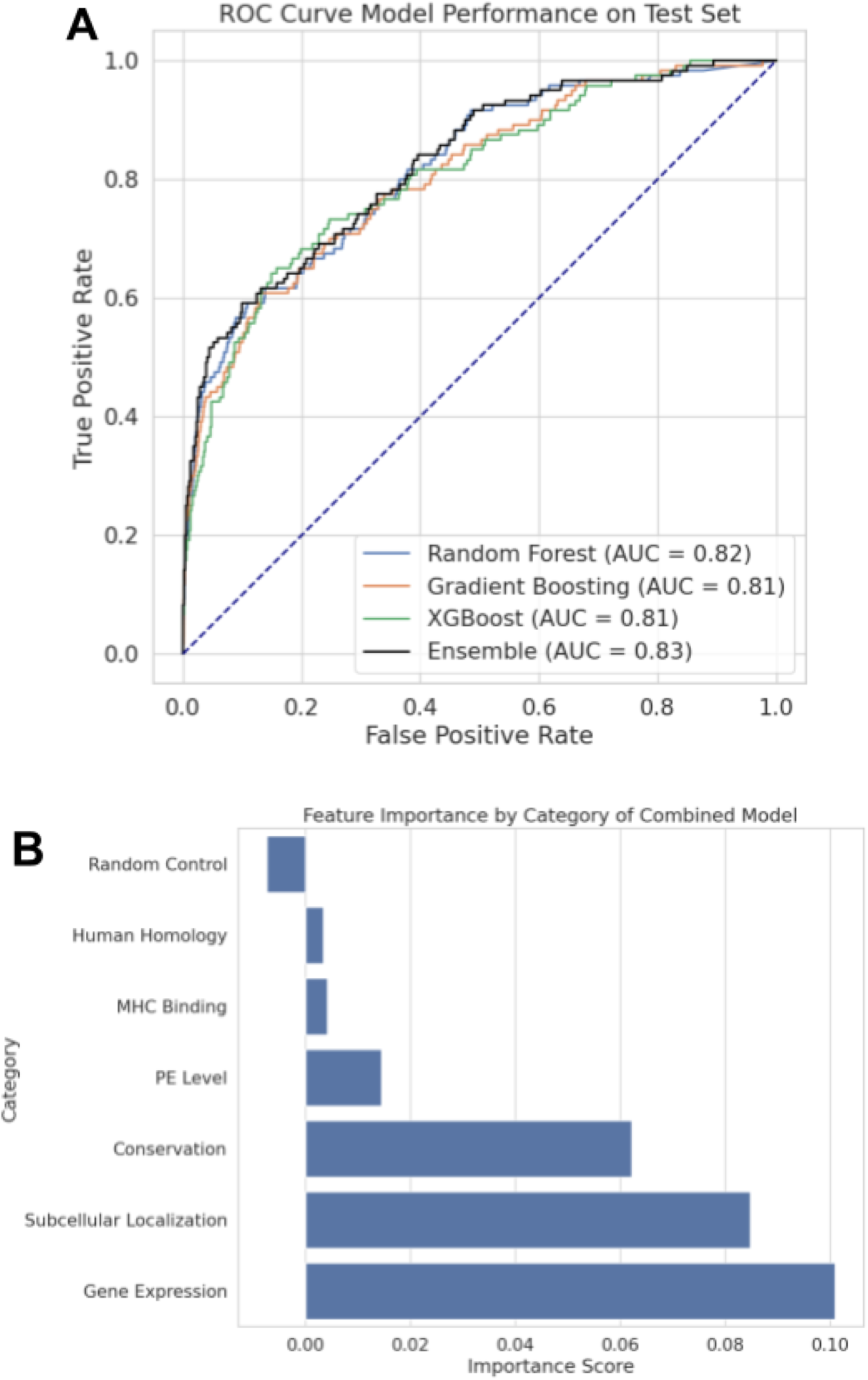
Performance of immunogenicity prediction by the trained models using the combined Mtb and *Bordetella pertussis data*. A) Performance evaluated as ROC-AUC for each of the three models and ensemble on the withheld set using the combined model trained on both Mtb and *B. pertussis* data. The ensemble model achieved the highest ROC-AUC of 0.83, outperforming each individual model. B) The feature importance by feature category. Gene expression had the greatest impact in the combined model, similar to the original Mtb model. However, conservation and subcellular localization had more impact in this combined model.

## 3. Methods

### 3.1 T cell Epitope Data

A total of 20,216 peptides, all 15-mers, derived from a proteome-wide screening of *Mycobacterium tuberculosis* (Mtb) using PBMCs from healthy IGRA+ participants (7) was used for training. The peptides were screened for recognition using an IFNγ ELISpot assay in 28 IGRA+ healthy individuals. They were derived from 21 Mtb genomes, parsing out all possible 15-mers from their protein sequences, with a total of 1,568,148 peptides. The initial peptide selection from this study was based on MHC class II binding predictions across 22 different HLA DR, DP, and DQ class II alleles using the IEDB consensus method.

For independent model validation on *Bordetella pertussis* peptides, we used an independent dataset consisting of 24,294 *Bordetella pertussis* peptides, which were screened in PBMCs from 20 participants for recognition (12). The dataset included the peptide sequence and the number of participant responses. All features were collected the same way as the IGRA+ dataset. To quantify the model performance on this dataset, we conducted ROC-AUC analysis at varying recognition thresholds, ranging from one to four participants.

Additionally, the trained ensemble model was used to predict the most immunogenic peptides in *Streptococcus pneumoniae*. All possible 15-mer residues (n=558,000) were derived from the *S. pneumoniae* (strain ATCC BAA-255/R6) proteome (UniProt proteome ID: UP000000586) and the corresponding features were collected for each peptide. A single peptide from each protein was selected based on the highest MHC-II binding prediction scores to maximize the influence of the other features in the model while still accounting for MHC-II binding.

### 3.2 Feature Collection

For model training, we collected multiple features that can be attributed to each Mtb peptide. These included MHC Class II binding predictions, RNA expression levels, conservation across Mtb strains and human proteins, subcellular localization, and protein existence scores. Each feature associated with the peptides was quantified, resulting in numeric values for all data points.

#### 3.2.1 MHC Class II Binding Predictions

The NetMHCIIpan 4.1 EL (eluted ligand) method was used to predict MHC binding across seven alleles (HLA-DRB1*03:01, HLA-DRB1*07:01, HLA-DRB1*15:01, HLA-DRB3*01:01, HLA-DRB3*02:02, HLA-DRB4*01:01, HLA-DRB5*01:01), which have been shown to represent a large HLA coverage of the human population. A “7-allele” score was calculated for each peptide by averaging the binding scores (ranging [0, 1]) for each allele.

#### 3.2.2 RNA Expression

RNA expression data for individual Mtb peptides were obtained from a GEO dataset: Series GSE229680 (14). This dataset includes high-throughput sequencing expression profiles of Mtb generated using the Illumina NextSeq 2000 platform. The gene expression values for the H37Rv Mtb strain cultured under normal conditions, as a control, were used. For each gene in the dataset, three control values for H37Rv, measured as transcripts per million (TPM) values, were averaged to obtain a single expression value per gene. Subsequently, using the gene symbols from which each peptide was derived, these averaged TPM values were mapped to their respective peptides. For *Bordetella pertussis* and *Streptococcus pneumoniae*, RNA expression data was sourced similarly from GEO Series GSE145049 (15) and GSE77748 (16), respectively, following the same approach for mapping averaged TPM values to their respective peptides.

#### 3.2.3 Conservation in Other Mtb Strains

PEPMatch (17), a peptide-matching tool, was used to determine the conservation of each peptide across various other Mtb strains. A total of 33 separate Mtb proteomes (supplementary material) were selected from UniProt (18) for their significance and non-redundancy. Each proteome was preprocessed by PEPMatch using k=3. Then, each peptide was searched across these proteomes, allowing up to three residue substitutions, corresponding to 80% homology for the 15-mer peptides. Four total conservation scores were calculated for each homology threshold (100%, 93%, 87%, and 80%) by totaling the number of matches at each threshold and dividing by the number of proteomes searched.

#### 3.2.4 Conservation in Human Host

Homology to the human host was also assessed using PEPMatch. The human proteome was downloaded from UniProt (ID: UP000005640), which included protein isoforms. Given the low conservation of these peptides in human proteins, each peptide was searched with a threshold of up to seven residue substitutions. The highest homology percentage for a given peptide match was then used as the score for this feature, which ranged from 100% to 53%. If no match was found for a peptide up to seven substitutions, the next homology percentage at eight substitutions (47%) was given.

#### 3.2.5 Subcellular Localization

Subcellular localization of peptides was assessed using PSORTb (11), a bioinformatics tool designed for predicting the localization of proteins within different cellular compartments. The provided Docker container was used to run the tool. Each protein sequence from which the Mtb peptides are derived was passed as the input to PSORTb. In addition, PSORTb requires a Gram staining indication, and Gram-negative was chosen for every Mtb protein. The tool categorizes proteins into four main subcellular locations: cytoplasm, cytoplasmic membrane, outer membrane, and extracellular space, and assigns a score ranging from 0 to 10 for each. A higher score indicates a greater likelihood that the protein, and by extension, the peptide derived from it, is localized in that particular cellular compartment.

#### 3.2.6 Protein Existence Levels

Each protein within the UniProt database is curated with a level ranging from 1 to 5 based on the evidence supporting its existence, from direct protein sequencing to bioinformatic predictions based on genomic data. Each Mtb peptide in our study was assigned the protein existence (PE) score of its corresponding protein from UniProt using its API and a custom Python script.

### 3.3 Model Training

Three machine learning methods, Random Forest, Gradient Boosting, and XGBoost, were used to train separate models on the Mtb peptides and their accompanying features. The Gradient Boosting and Random Forest models were trained using the scikit-learn library, and the XGBoost model was trained separately with its independent Python bindings. Hyperparameter tuning using grid search with stratified 5-fold cross-validation was used for each model. The best estimator from each grid search was taken for the final model from each method. For Random Forest, hyperparameters included ’n_estimators’ [100, 200, 300], ’max_depth’ [None, 10, 20], ’min_samples_split’ [2, 5, 10], ’min_samples_leaf’ [1, 2, 4], and ’max_features’ [’auto’, ’sqrt’]. For Gradient Boosting, the grid search covered ’n_estimators’ [100, 200, 300], ’learning_rate’ [0.01, 0.05, 0.1, 0.2], ’max_depth’ [3, 5, 7], and ’subsample’ [0.5, 0.75, 1.0]. The XGBoost model’s hyperparameters included ’n_estimators’ [100, 200, 300], ’learning_rate’ [0.01, 0.1, 0.2], ’max_depth’ [3, 5, 7], ’subsample’ [0.8, 1], ’colsample_bytree’ [0.8, 1], ’lambda’ [1, 1.5, 2], ’alpha’ [0, 0.5, 1], and ’scale_pos_weight’ [1, ratio of negative to positive samples]. After training was performed for all three methods, an ensemble model was created by averaging the probabilities from each model prediction.

### 3.4 Model Evaluation

The models were evaluated on the withheld test data using ROC-AUC values (19) as well as AUC0.1, which is the area under the ROC curve from 0-10% false positive rate. These measures were also calculated for the ensemble model, combining the probabilities of the trained models. The evaluation compared the effectiveness in predicting peptide immunogenicity using the recognition of different numbers of total participants.

### 3.5 Feature Importance

To assess the contribution of each feature to the model performance, we employed importance-by-permutation provided by scikit-learn. The scikit-learn “inspection” submodule includes a permutation function for scoring each method. The permutation importance was calculated for each of the Random Forest, Gradient Boosting, and XGBoost models on the withheld test data, with 10 repeats, using the AUC values as the scoring metric for the model’s decrease in performance. In addition, a control feature was added that assigned a random value between 0 and 1 to each peptide, and this was included during training to determine its ranking amongst all features.

### 3.6 *Ex vivo* validation of the Ensemble Model using the *Streptococcus pneumoniae* Proteome

The peptides with the highest and lowest ensemble probability scores were classified as predicted to be immunogenic (n=130) and non-immunogenic (n=130). Peptides were purchased from TC Peptide Lab (San Diego) as crude material on a 1 mg scale. Peptides were combined into 26 pools of 10 peptides each (13 pools with high probability scores, 13 pools with low probability scores), along with a “megapool” consisting of all 260 peptides. The megapool (20) was constructed by pooling the peptides, followed by lyophilization to increase the stock concentration of the peptides resuspended in DMSO. The 10-peptide pools (2 mg/ml) and megapool (1 mg/ml) were stored in aliquots at -20°C until the point of use.

#### 3.6.1 Participant Selection and PBMC Preparation

All participants (n=20) provided written informed consent for participation in the study, and ethical approval was obtained from the institutional review boards at La Jolla Institute for Immunology (LJI IRB VD-071/VD-101). All individuals were >18 years old, free from any acute infection, and recruited in San Diego, California. Peripheral blood mononuclear cells (PBMCs) were isolated from whole blood by density gradient centrifugation with Ficoll (VWR International) according to the manufacturer’s instructions. The cells were cryopreserved in liquid nitrogen and suspended in fetal bovine serum (FBS) containing 10% dimethyl sulfoxide (DMSO).

Each PBMC cryovial was thawed at 37 °C for 2 min, and cells were transferred to medium (RPMI 1640 with L-glutamine and 25 mM HEPES; VWR International), supplemented with 5% human AB serum (Gemini Bio), 1% penicillin-streptomycin (Gemini Bioproducts), 1% glutamax (Gibco) and 20 U/ml benzonase nuclease (Fisher Scientific). Cells were centrifuged and resuspended in RPMI medium to determine cell concentration using trypan blue.

#### 3.7.2 Peptide Screening and Deconvolution

The synthesized peptides were screened for recognition with IFNγ Fluorospot assays. Participant responses were determined based on the number of IFNγ spot-forming cells (SFC) following stimulation with peptide pools or individual peptides and controls. Peptides were tested in 10-peptide pools, along with the megapool consisting of all 260 peptides. All 10-peptide pools with significant IFNγ response were deconvoluted to determine individual peptide responses. Assays were performed in triplicates, except for the DMSO negative control, which was tested in six wells. The background was calculated by averaging the six DMSO wells by participant. The SFC was calculated by averaging the triplicate wells for each stimulation condition. Responses were considered positive if they met the following criteria: 1) more than 20 SFC per 10^6^ PBMC, 2) a greater than 2-fold increase above background, and 3) statistical significance (p < 0.05) based on an independent t-test and Poisson distribution test when comparing stimuli to negative control. A maximum cutoff of 500 SFC/2×10^5^ was utilized. Any responses that did not meet the 3 criteria for positivity were plotted as 0 SFC/2×10^5^ cells to highlight IFNγ responses that were significantly different compared to the negative control. The magnitude of IFNγ SFC per well (2×10^5^ cells) of immunogenic versus non-immunogenic peptides was compared with an independent one-tailed t-test. The response frequency was calculated by dividing the number of participants with a response by the total number of participants screened.

Plates were coated overnight at 4°C with an antibody mixture containing mouse anti-human IFNγ (Mabtech, 1-D1K). The coated plates were then prepped with stimuli as follows: megapool at 2 μg/ml, individual pools at 5 μg/ml, PHA at 10 μg/ml (positive control), and media containing 0.2% DMSO (negative control). The stimuli were then combined with 2×10^5^ PBMCs per well for 20-24 hours at 37°C in a humidified CO_2_ incubator. After incubation, the cells were removed and the wells were washed with PBS/0.05% Tween with an automatic plate washer. An antibody mixture containing anti-IFNγ (7-B6-1-FS-BAM) (Mabtech) was prepared in PBS with 0.1% BSA and incubated for 2 hours at room temperature. The plates were then washed and incubated with anti-BAM-490 (Mabtech) for 1 hour at room temperature. A final wash and incubation with a fluorescence enhancer (Mabtech) for 15 minutes was performed and the fluorescent spots were counted with the IRIS Fluorospot reader (Mabtech).

### 3.7 Combined Model Training on Mtb and B. pertussis Data

We combined the peptide datasets from Mtb and *Bordetella pertussis* for model training. The combined dataset comprised 44,510 peptides, with 20,216 peptides from Mtb and 24,294 peptides from B. pertussis. Positive peptides were defined based on recognition by at least two participants in their respective datasets, resulting in 144 positive peptides from Mtb and 412 positive peptides from B. pertussis, for a total of 556 positive peptides. We retrained the Random Forest, Gradient Boosting, and XGBoost algorithms on the combined dataset using the same hyperparameter tuning and stratified 5-fold cross-validation procedures described in Section 3.3. The ensemble model was constructed by averaging the predicted probabilities from each of the three individual models. Model performance was evaluated as described in Section 3.4. Feature importance was performed as described in Section 3.5.

## 4. Discussion / Conclusion

This study demonstrates the effectiveness of machine learning models in predicting immunogenic CD4+ T cell epitopes across multiple bacteria causing airway infections, with a particular focus on Mtb, *Bordetella pertussis*, and *Streptococcus pneumoniae*. Integrating multiple biological features, including MHC class II binding predictions, RNA expression levels, conservation scores, and subcellular localization, allowed for the developing of robust predictive models. Among these features, gene expression levels and conservation emerged as the most critical factors influencing epitope immunogenicity. These findings highlight the importance of both the level of antigen expression within the pathogen and the shared homology across Mtb strains in determining the likelihood of eliciting a T cell response. The low feature importance of MHC binding predictions, which would normally be an important factor, is likely due to the datasets being pre-filtered for this criterion. Our ensemble model, combining the predictions from Random Forest, Gradient Boosting, and XGBoost algorithms, showed superior performance in identifying epitopes with high immunogenic potential. This approach not only streamlines the epitope selection process, reducing the need for extensive peptide synthesis and experimental validation, but also provides insights into the important biological features of Mtb infection.

The validation of our models using data from *Bordetella pertussis* peptides highlights the model’s generalizability across different pathogens and disease states. The progressive increase in predictive accuracy observed with higher thresholds for the number of participants recognizing each epitope in the *Bordetella pertussis* validations indicates that our model is particularly effective in identifying epitopes that elicit broad immune recognition. This is crucial for vaccine design, as epitope targets recognized by a larger proportion of the population are beneficial for inducing protective immunity in a population. Additionally, the ability of the model to distinguish between immunogenic and non-immunogenic peptides in *Streptococcus pneumoniae*, as evidenced by the ex vivo IFNγ FluoroSpot assays, further underscores the practical utility of our approach in guiding experimental efforts. This approach can be applied to large-scale epitope discovery efforts, particularly for pathogens with few or no known CD4+ T cell targets.

The features for our model were chosen due to their applicability across a wide range of pathogens, particularly because they are relatively easy to obtain and do not require extensive pathogen-specific data. However, our study has limitations. While our model effectively predicts epitopes based on the features included, there are likely several, if not many, other relevant factors influencing immunogenicity that were not captured in this study. Also, the validation efforts included diverse bacterial pathogens but did not include viral pathogens, which also can cause pneumonia. Extending the model to more pathogens and pathogen types could provide further insights into its applicability across a broader range of infectious agents. Finally, while the model showed high accuracy in predicting immunogenicity, the challenge of producing effective vaccine targets is much more complicated in vivo, and our study does not account for this.

We also set out to combine the Mtb and *Bordetella pertussis* data by training a new model using the same methods. Although we did not conduct experimental validation for this combined model within the scope of this study, and the AUC values are lower than the original Mtb model, we believe by encompassing multiple bacterial species, the model can capture a broader candidate of epitopes from pathogens with limited immunological data. We intend to make this combined model publicly available, providing a valuable resource for the research community.

Predicting CD4+ T cell epitopes in Mtb and other pneumonia-causing bacteria using machine learning models offers a promising approach to identifying the most likely immunogenic peptides, reducing the need to synthesize and test large numbers of peptides, which can be both time-consuming and costly. Machine learning models can integrate many biological features to predict which epitopes are likely to elicit strong immune responses. This peptide prediction model has huge applications for vaccine discovery and diagnostic development since it is capable of rapidly identifying the most immunogenic bacterial epitopes. This study leveraged computational approaches to predict immunogenic CD4+ T cell epitopes across an array of different respiratory diseases, highlighting the diverse applicability of this methodology.

In conclusion, our study demonstrates that machine learning models can significantly enhance the efficiency and accuracy of CD4+ T cell epitope prediction, offering a powerful tool for vaccine development and immunological research. By reducing the need for large-scale peptide synthesis and experimental screening, this approach has the potential to accelerate the identification of immunogenic epitopes and contribute to the development of more effective vaccines against a range of bacterial pathogens. Future studies should focus on further refining these models, exploring their application to other pathogens, and validating their predictions in clinical and experimental settings.

## Author Contributions

DM collected the data, wrote the code to train and evaluate the models, and authored the bioinformatics portion of the manuscript. HB performed the experimental work, analyzed the results, and authored the experimental portion of the manuscript. EK and MM aided in feature selection and model tuning. SP provided the Mtb dataset. RA provided the *Bordetella pertussis* dataset. AS and CA aided in the design of the work and provided feedback. BP designed and supervised both the bioinformatics and experimental work.

## Supporting information

mtb-proteomes

